# Cortical auditory distance representation based on direct-to-reverberant energy ratio

**DOI:** 10.1101/769380

**Authors:** Norbert Kopco, Keerthi Kumar Doreswamy, Samantha Huang, Stephanie Rossi, Jyrki Ahveninen

## Abstract

Auditory distance perception and its neuronal mechanisms are poorly understood, mainly because 1) it is difficult to separate distance processing from intensity processing, 2) multiple intensity-independent distance cues are often available, and 3) the cues are combined in a context-dependent way. A recent fMRI study identified human auditory cortical area representing intensity-independent distance for sources presented along the interaural axis (Kopco et al., PNAS, 109, 11019-11024). For these sources, two intensity-independent cues are available, *interaural level difference* (ILD) and *direct-to-reverberant energy ratio* (DRR). Thus, the observed activations may have been contributed by not only distance-related, but also direction-encoding neuron populations sensitive to ILD. Here, the paradigm from the previous study was used to examine DRR-based distance representation for sounds originating in front of the listener, where ILD is not available. In a virtual environment, we performed behavioral and fMRI experiments, combined with computational analyses to identify the neural representation of distance based on DRR. The stimuli varied in distance (15-100 cm) while their received intensity was varied randomly and independently of distance. Behavioral performance showed that intensity-independent distance discrimination is accurate for frontal stimuli, even though it is worse than for lateral stimuli. fMRI activations for sounds varying in frontal distance, as compared to varying only in intensity, increased bilaterally in the posterior banks of Heschl's gyri, the planum temporale, and posterior superior temporal gyrus regions. Taken together, these results suggest that posterior human auditory cortex areas contain neuron populations that are sensitive to distance independent of intensity and of binaural cues relevant for directional hearing.

**Highlights:**

- Posterior auditory cortices (AC) are sensitive to frontally presented distance cues
- These effects are independent of intensity- and direction-related binaural cues
- fMRI activations to frontal distance cues are found in the right and left AC
- The frontal reverberation-related auditory distance cues are behaviorally relevant

## Introduction

Information about distance of objects that surround us in the environment is often important. The auditory modality is special in that it provides such information even for objects that are occluded or behind the listener (Brungart and Simpson, 2002b; Genzel et al., 2018; Kolarik et al., 2016; Maier et al., 2004; Neuhoff, 1998; Shinn-Cunningham et al., 2001; Zahorik et al., 2005). A reliable cue for auditory distance is the overall received stimulus level, which dominates distance perception of familiar objects and of looming vs. receeding sound sources (Ghazanfar et al., 2002; Hall and Moore, 2003; Maier and Ghazanfar, 2007; Seifritz et al., 2002). However, in many situations, the emitted sound level is varying or unknown. In such cases, auditory distance perception can only rely on intensity-independent cues.

Previous psychoacoustic studies showed that distance perception is possible without the overall intensity cue, especially when the sources are in the peripersonal space (up to 1 – 2 meters from the listener), in which the listener can interact with the objects producing the sounds (Kolarik et al., 2016; Shinn-Cunningham et al., 2000). Nearby intensity-independent distance judgements might be particularly relevant in social situations in which the listener concentrates on a nearby speaker in a conversation (Brungart and Simpson, 2002a; Westermann and Buchholz, 2015) and when the emitted level of surrounding sounds naturally fluctuates (Bronkhorst and Houtgast, 1999; Maier et al., 2004; Shinn-Cunningham et al., 2000; Zahorik et al., 2005). Two dominant intensity-independent distance cues were previously identified. First, in reverberant environments, distance judgements can be made by comparing the sound received by the ears directly from the source vs. its reflections off the walls (the direct-to-reverberant energy ratio; DRR) (Hartmann, 1983; Mershon and King, 1975; Zahorik, 2002). Second, for nearby sources located off the midline, intensity-independent distance judgements can be based the interaural level difference (ILD) (Brungart, 1999; Shinn-Cunningham et al., 2005). While the relative contribution of these two cues to intensity-independent distance perception is currently not known, it is likely that it is context-dependent, varying with target azimuth, distance, as well as with the availability of the reverberation cue. For example, based on acoustic analysis and modeling of behavioral performance in experiments in which azimuth as well as distance was varied, Kopco *et al.* (2011) suggested DRR is the main distance cue in reverberation. However, a similar analysis performed on sources varying in distance for a single direction, along the interaural axis, suggested that both ILD and DRR cues were used (Kopco et al 2012). Further studies are therefore needed to clarify how the intensity-independent distance cues are combined in various contexts.

Although the fine-grained functional arrangement of human auditory cortices (AC) is not fully understood, an abundance of human neuroimaging evidence exists of their broader anatomical subdivisions and functional pathways. According to these studies (Ahveninen et al., 2006; Rauschecker, 1997, 1998; Rauschecker and Tian, 2000; Rauschecker et al., 1995), auditory-spatial feature changes activate most strongly the posterior non-primary AC areas (i.e., the “where” pathway), including the planum temporale (PT) and posterior superior temporal gyrus (STG). In contrast, attributes related to the sound-source identity could be processed predominantly in the more anterior “what” pathway. Consistent with the what-where dichotomy, it has been well documented that posterior non-primary ACs are strongly activated by horizontal sound direction changes (Ahveninen et al., 2006; Brunetti et al., 2005; Deouell et al., 2007; Tata and Ward, 2005) and movement (Krumbholz et al., 2005; Warren et al., 2002). However, neuronal representations of distance have been studied much less intensively. Our previous fMRI study (Kopco et al., 2012) provided evidence of neuron populations sensitive to intensity-independent auditory distance cues in these spatially-sensitive AC areas as well. However, this evidence was obtained for sources simulated to originate at the side of the head that varied in distance along the interaural axis, for which both DRR and ILD cues are available. It is thus possible that the findings are an epiphenomenon of activations of direction-encoding neurons that are sensitive to ILD (Imig et al., 1990; Johnson and Hautus, 2010; Lehmann et al., 2007; Tardif et al., 2006; Zimmer et al., 2006), which have been later shown to activate areas overlapping with the putative distance representations (Higgins et al., 2017; Stecker et al., 2015). Further studies are, therefore, needed to verify the existence of auditory cortex distance representations that do not involve cues shared with directional hearing.

Here, one behavioral and one imaging experiments were performed in a virtual auditory environment. The experiments examined intensity-independent distance perception for *frontal* sources, for which no ILD cue is available and distance judgements are expected to be naturally based on DRR. In the behavioral experiment we verified that intensity-independent distance perception is possible for the frontal sources in reverberation, and that performance for frontal sources is worse than for lateral sources for which both ILD and DRR cues are available. Then, in the imaging experiment, we used a sparse sampling adaptation fMRI paradigm to compare responses to frontal sources varying in distance vs. frontal sources at a fixed distance and varying only in intensity, to identify the AC area encoding intensity-independent DRR-based distance information.

## Methods

### Subjects

Fourteen subjects (4 females, ages 20 – 41 years) with normal hearing (audiometric thresholds within 20 dB HL) participated in the behavioral experiment. The behavioral experimental protocol was approved by the P. J. Šafárik University (UPJŠ) Ethical Committee. A separate sample of 12 right-handed individuals (4 females, ages 22 – 55 years) with self-reported normal hearing participated in the imaging experiment. The protocol of the imaging experiment was approved by the Partners Human Research Committee, the Institutional Review Board (IRB) of the MGH. All subjects gave a written informed consent to participate in the study. fMRI data of one imaging subject were excluded due to excessive head motion during the experiment.

### Stimuli

The auditory distance stimuli were simulated using a single set of non-individualized binaural room impulse responses (BRIR) measured on a listener that did not participate in this study, using procedures and devices that were, unless specified otherwise, identical to our previous studies (Kopco et al., 2012; Shinn-Cunningham et al., 2005). We measured the BRIRs in a small carpeted classroom (3.4 m × 3.6 m × 2.9 m height) with hard walls and acoustic-tile ceiling, using a surface-mount cube speaker (Bose FreeSpace 3 Series II, Bose, Framingham, MA). The room reverberation times, T_60_, in octave bands centered at 500, 1000, 2000, and 4000 Hz ranged from 480 to 610 ms. Miniature microphones (Knowles FG-3329c, Knowles Electronics, Itasca, IL) were placed at the blocked entrances of the listener’s ear canals and the loudspeaker was set to face the listener at various distances (15, 19, 25, 38, 50, 75, or 100 cm) from the center of the listener’s head at the level of the listener’s ears (Fig. 1A). The recordings were made for two directions, either in front of the listener or on the left-hand side along the interaural axis. Analogously to our previous study (Kopco et al., 2011), a set of 50 independent noise burst tokens that consisted of 300-ms white-noise samples filtered at 100–8000 Hz was then convolved with each of the BRIRs to create standard stimuli for each source distance and direction. An otherwise identical set of 150-ms deviant stimuli was also generated for the fMRI experiments. For each experimental trial, either two (behavioral experiment, Fig. 1C) or 14 (imaging experiment, Fig. 1D) noise bursts were randomly selected, scaled depending on the normalization scheme used, and placed in a series with a fixed stimulus onset asynchrony (SOA), to create the stimulus sequence. Finally, the fMRI stimuli were filtered to compensate for the headphone transfer functions.

**Figure 1.**
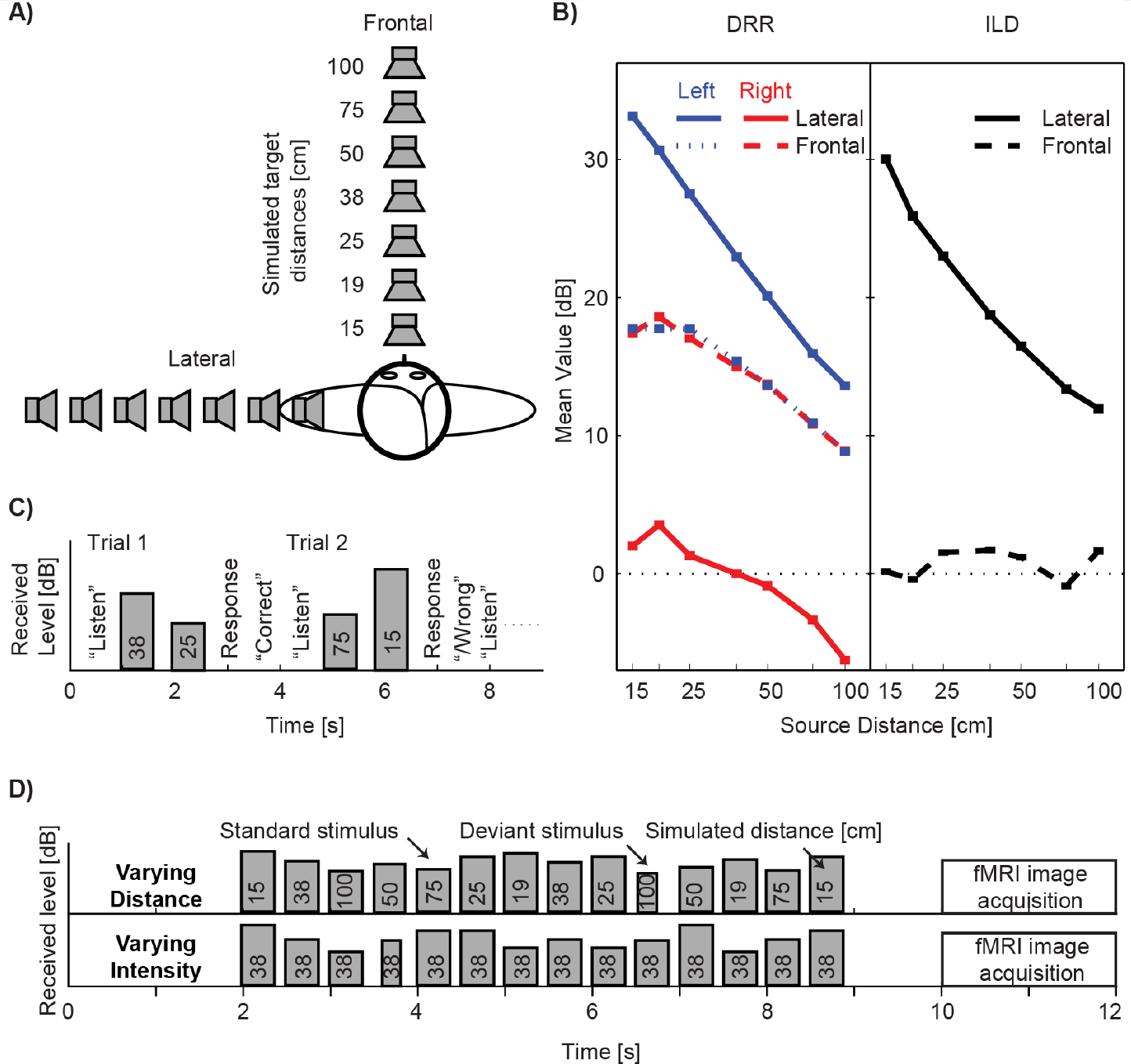
Experimental design. (**A**) Simulated source locations. Note that only the frontal locations were used in the fMRI experiment. (**B**) Mean values of DRR (Left) and ILD (Right) as a function of distance for the frontal and lateral stimuli used in this study. Values were computed for the whole broadband stimulus separately for each combination of distance and direction. Error bars represent SDs across random noise tokens used as stimulus. (**C**) Timing of events during trials in the behavioral experiment: The instruction “listen” appeared on the screen, followed by presentation of two stimuli from different distances. Listeners responded by indicating whether the second stimulus sounded more or less distant than the first stimulus. On-screen feedback was provided. Stimulus bar height shows that presentation intensity was randomly roved for each stimulus so that received intensity could not be used as a cue in the distance discrimination task. (**D**) Timing of stimuli and of image acquisition during one imaging trial in the fMRI experiment, shown separately for the two stimulus conditions used in the experiment. Height of the stimulus bars corresponds to the received stimulus intensity at the listener’s ears. In the varying distance condition, the stimulus distance changed randomly, whereas the stimulus presentation intensity was fixed (thus, both the perceived distance and intensity varied). In the varying intensity condition, the stimulus distance was fixed, whereas the received intensity varied over the same range as in the varying distance condition. In these conditions, the listener’s task was to detect deviant stimuli that were shorter than the standard stimuli. No feedback was provided.

All stimuli were pre-generated offline at a sampling rate of 44.1 kHz. Average received level was 65 dBA (measured at the left ear of a KEMAR manikin equipped with the DB-100 Zwislocki Coupler and the Etymotic Research ER-11 microphones). The stimuli used in the experiments mainly differed by the overall stimulus intensity normalization used. In the behavioral experiment, each noise burst was normalized so that its overall intensity received at the left ear (which was closer to the lateral simulated sources) was fixed and then randomly roved over a 12-dB range so that the monaural overall intensity distance cue was eliminated at both the left and right ears for both frontal and lateral stimuli (Fig. 1C). In the imaging experiment, two types of stimuli were used. The varying distance stimuli were normalized such that the presentation intensity was fixed (i.e., the overall intensity cue was present in these stimuli because the received intensity for the near sources was higher than the received intensity for the far sources). The varying intensity stimuli were simulated from the fixed distance of 38 cm and their presentation intensity was varied such that the received intensity at the ears varied across the same range as for the varying distance stimuli.

### Behavioral experiment

The behavioral experiment was performed in the Perception and Cognition Lab at UPJŠ. The subjects were seated in a double-walled sound-proof booth in front of an LCD display and a keyboard connected to a control computer which ran a Matlab (Mathworks) script controlling the experiment. The pre-generated stimuli were played through Fireface 800 sound processor (RME) and Etymotic Research ER-1 insert earphones. Each subject performed one 1-hour-long session consisting of 3 runs with frontal stimuli and 3 with lateral stimuli in a random order (stimulus direction was fixed within a run; additional 3 experimental runs were performed in each session, data of which are not included in this analysis). Each run consisted of 84 trials, corresponding to 4 repetitions of 21 randomly ordered trials (one trial for each combination of 2 out of the 7 distances). Each trial started with the word “Listen” appearing on the computer screen, followed after 200 ms by two noise tokens simulated from two different distances with a 1000 ms SOA. The subject was asked to indicate whether the second sound source was closer or farther away than the first source by pressing one of two keys on the keyboard. Feedback was provided after the response. The experiment was self-paced and the total duration of one trial was, on average, approximately 5 s.

### Imaging experiment

The imaging experiment was performed at the Martinos Center for Biomedical Imaging at MGH. The subjects participated in one 2-hour-long session inside a research 3T fMRI magnet. The control computer, running a Presentation (Neurobehavioral Systems) experimental script, presented the sounds through the Fireface 400 sound processor (RME), Pyle Pro PCA1 amplifier, and Sensimetrics S14 (Sensimetrics, Gloucester, MA) MRI-compatible headphones. Responses were collected via MRI-compatible five-key universal serial bus (USB) keyboard. A video projector was attached to the control computer and projected the instructions to the subject in the scanner. A single run of 96 trials was performed. Each trial consisted of a 10-s stimulus presentation during which the scanner was silent, followed by 2 seconds of fMRI image acquisition (Fig. 1D). Trials with two types of stimuli, varying in distance or varying in intensity, were randomly interleaved. Each stimulus consisted of a sequence of 14 noise bursts with SOA of 500 ms. The varying-distance sequences contained two noise bursts for each of the seven distances, ordered pseudo-randomly such that each distance was present at least once before the second occurrence of any of the distances. The varying-intensity sequences contained 14 bursts simulated from the fixed distance of 38 cm and only varying in their intensity. In 50% of the sequences, one randomly chosen burst out of the 14 bursts was replaced by a 150-ms deviant. The listener’s task during the fMRI session was to detect these short-duration deviants and to respond by a button press whenever they heard one.

## Data acquisition

Whole-head fMRI was acquired at 3T using a 32-channel coil (Siemens TimTrio, Erlagen, Germany). To circumvent response contamination by scanner noise, we used a sparse-sampling gradient-echo blood oxygen level dependent (BOLD) sequence (TR/TE = 12,000/30 ms, 9.82 s silent period between acquisitions, flip angle 90°, FOV 192 mm) with 36 axial slices aligned along the anterior-posterior commissure line (3-mm slices, 0.75-mm gap, 3×3 mm^2^ in-plane resolution). The coolant pump was switched off during the acquisitions. T1-weighted anatomical images were obtained using a multi-echo MPRAGE pulse sequence (TR=2510 ms; 4 echoes with TEs=1.64 ms, 3.5 ms, 5.36 ms, 7.22 ms; 176 sagittal slices with 1×1×1 mm^3^ voxels, 256×256 mm^2^ matrix; flip angle = 7°) for combining anatomical and functional data.

## Data analysis

### Behavioral Data

In the distance discrimination behavioral experiment, the proportion of correct responses was analyzed for each distance pair, direction, repeat and subject. We then computed the across-subject means and standard errors of the mean of the data averaged across direction and repeat. To analyze the dependence of discrimination performance on source-pair distance, a linear regression model was fitted to data grouped by the number distance intervals between source pairs. A statistical test was performed to determine whether the slope was significantly different from 0.

In the duration discrimination task performed during the imaging experiment, the response was classified as a hit (correct detection) if it occurred within 2.5 s after the deviant onset. We determined the hit rates (HR) and reaction times (RT) to correctly detected targets and compared the group averages of these measures across the two stimulus conditions. Statistical comparisons of the behavioral data were done using repeated measures ANOVAs (CLEAVE, http://www.ebire.org/hcnlab/software/cleave.html). Percent correct data were rau-transformed before submitting them to ANOVA.

### fMRI Data

Cortical surface reconstructions and standard-space co-registrations of each subject’s anatomical data (Dale et al., 1999) and the functional data analyses were conducted using Freesurfer 5.3. Individual functional volumes were motion corrected, coregistered with each subject’s structural MRI, intensity normalized, resampled into standard cortical surface space (Fischl et al., 1999a; Fischl et al., 1999b), smoothed using a 2-dimensional Gaussian kernel with an FWHM of 5 mm, and entered into a general-linear model (GLM) with the task conditions as explanatory variables. A random-effects inverse-variance weighted least-squares (WLS) GLM was then conducted at the group level. A volumetric statistical analysis was conducted to enhance the comparability of our results to previous studies. In this case, each subjects native voxel data were smoothed with a 3-dimensional (3D) kernel with a 5 mm FWHM. The resulting contrast effect size estimates were coregistered to the Montreal Neurological Institute (MNI) 305 standard brain representation (2 × 2 × 2 mm^3^ resolution) for a volumetric random-effects WLS-GLM. In the surface-based analyses, multiple comparisons were controlled for using a cluster-based Monte Carlo simulation test with 5,000 iterations, with an initial cluster-forming threshold *p*<0.01 (two tails). In the 3D group analysis, multiple comparison problems were handled based on the theory of the global random fields (GRF). All surface-based results are rendered in the Freesurfer “fsaverage” standard subject cortex representation. The results of the volumetric group analyses are shown in a 3D rendering produced by using ITK-SNAP (Yushkevich et al., 2006) and ParaView (Ayachit, 2005), as described in (Madan, 2015).

Finally, we also conducted an *a priori* region-of-interest (ROI) analysis, which was based on the cortical surface-based labels that were used in our previous study (Kopco et al., 2012). One ROI was defined in each hemisphere by combining two anatomical FreeSurfer standard-space labels encompassing PT and posterior aspect of STG. To test our hypothesis, we quantified the activations as contrast effect sizes converted to percent-signal changes (PSC), which were normalized by the variance of PSC in each surface vertex location of these labels, before computing the ROI averages in each subject. The resulting activation magnitude measures were then analyzed at the group level using a non-parametric randomization test, which adjusts the p-values of each variable for multiple comparisons (Blair and Karniski, 1993; Manly, 1997).

## Results

### Behavioral Experiment

Fig. 2 plots the results of the behavioral distance discrimination experiment. It shows percent correct distance discrimination as a function of separation between speakers (for individual speaker pairs, as well as averaged across speaker pairs), separately for the frontal and lateral directions. Specifically, since the target locations were distributed approximately uniformly on a logarithmic scale, the distance between speakers in a pair is expressed as a number of intervals between speakers, such that a separation of one interval corresponds to a pair of neighboring speakers (*e.g.*, 15-19 cm, 19-25 cm, etc., see Fig. 1A), two intervals correspond to speaker pairs (e.g., 15-25 cm, 19-35 cm), etc. Thin lines at each separation show the data for each individual speaker pair as a function of their distance (e.g., at 1-interval separation there are 6 pairs, 15-19, 19-25, 25-35 cm etc.). No significant effects were observed as a function of speaker pair location (regression slopes were fitted to data for each source-pair group; then, two-tailed Student t test with Bonferroni correction performed on the fitted slopes found no significant deviation from 0; P > 0.1). So, the data for each interval were averaged across speaker-pair location (across 6 speaker pairs for 1 interval, across 5 for 2 intervals, etc.) and plotted using thick lines. The results in Fig. 2 show that, as expected, the listener’s accuracy improved with increasing distance difference between the two simulated sound sources (both lines grow with increasing number of intervals). Moreover, the lateral distance discrimination was much more accurate than the frontal discrimination (solid line always falls above the corresponding dashed line). Supporting these observations, a repeated measures ANOVA with the factors of separation (6 levels) and direction (frontal X lateral) performed on RAU-transformed percent correct data found a significant main effect of separation (*F*_5,65_ = 34.59, *p* < 0.0001) and direction (*F*_1,13_ = 50.16, *p* < 0.0001), but no significant interraction (*F*_5,65_ = 1.62, *p* > 0.16).

**Figure 2.**
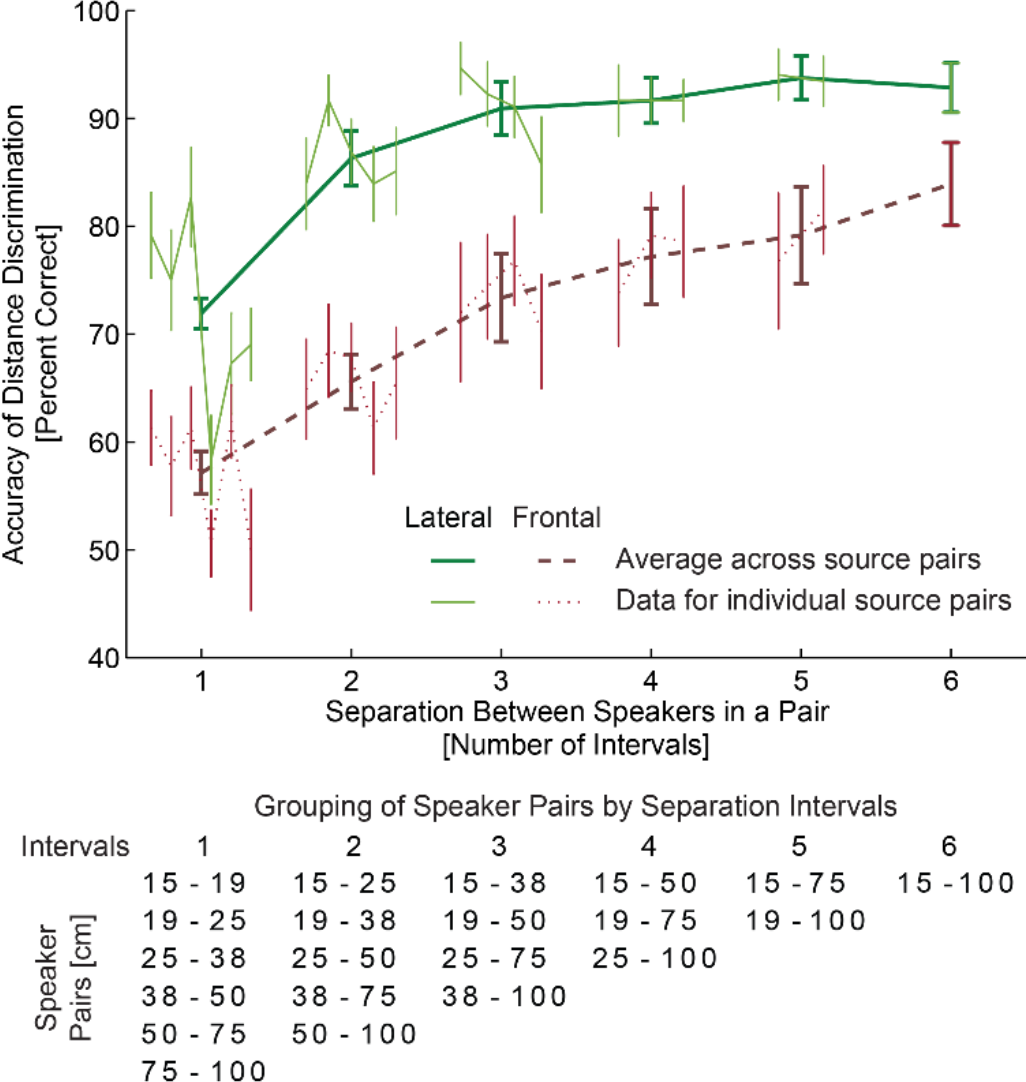
Behavioral distance discrimination responses. The thick lines show across-subject average accuracy collapsed across simulated source pairs separated by the same number of unit log-distance intervals, which are specified in the table below the graph (see also Fig. 1A), separately for the frontal and lateral directions (green vs brown). The accuracy improved as the simulated source separation (number of intervals) increased, reflecting the robustness of our virtual 3D stimuli. Thin lines show across-subject average performance separately for each source-distance pair. Each line represents a grouping on the basis of the number of intervals between sources within the pair (table below lists in each column the source pairs that are separated by the same number of intervals. No systematic upward or downward trend is visible in performance across source pairs within each group. Overall performance was better when sounds were simulated laterally along the inter-aural axis (thick solid green line), with both ILD and DRR cues available. The subjects were able to discriminate the distances from the frontal direction (thick dashed brown line). The error bars represent standard errors of the mean.

Importantly for the current fMRI experiment, the listeners were able to perform intensity-independent distance discrimination for the frontal sources even at the smallest separation between the speakers, confirming that our virtual auditory environment was robust and that the listeners were able to extract intensity-independent distance cues from the stimuli. In fact, it can be expected that the distance percepts were even more robust in the fMRI experiment, as the overall level cue was also present there.

### fMRI experiment

To assure that the brain activations measured in the fMRI experiment are not contaminated by fluctuations in attention and alertness during the scanning, subjects were asked to detect occasional changes in sound duration that occurred independently of distance or intensity during both distance and intensity trials. Analyses of hit rates (HR) and reaction times (RT) were performed. One subject did not perform the task correctly due to misunderstanding the task and another subject’s data were lost due to malfunction of the response device. For the remaining 10 subjects, the task difficulty was similar across the different stimulus types (across-subject average HR of 93.3% and 90.8% and RT of 1164 and 1181 ms, respectively, for the varying distance and varying intensity conditions), suggesting that any fMRI activation differences across conditions cannot be attributed to differences in task difficulty or subject’s vigilance. Confirming these observations, repeated measures ANOVAs performed on the HRs and RTs found no significant differences (HR: *F*_1,9_ = 6.75, *p* > 0.31; RT: *F*_1,9_ = 4.5, *p* > 0.19).

To localize intensity-independent and ILD-independent distance representations, we compared auditory cortex areas activated during distance vs. intensity changes (Fig. 3). Specifically, we compared fMRI areas activated to sounds simulated from various distances (Varying Frontal Distance) *vs*. sounds varying in intensity only and presented at a fixed distance of 38 cm (Varying Intensity). In support of our hypothesis, the contrast between the Varying Frontal Distance and Varying Intensity conditions revealed a significant difference in both left and right posterior non-primary AC areas (cluster-based MCMC simulation test, two-tail *p*<0.01). In the “fsaverage” brain surface representation, these differences extended from the posterior bank of Heschl’s gyrus (HG) to the posterior aspects of superior temporal gyurs (STG) and planum temporale (PT) (Fig. 3).

**Figure 3.**
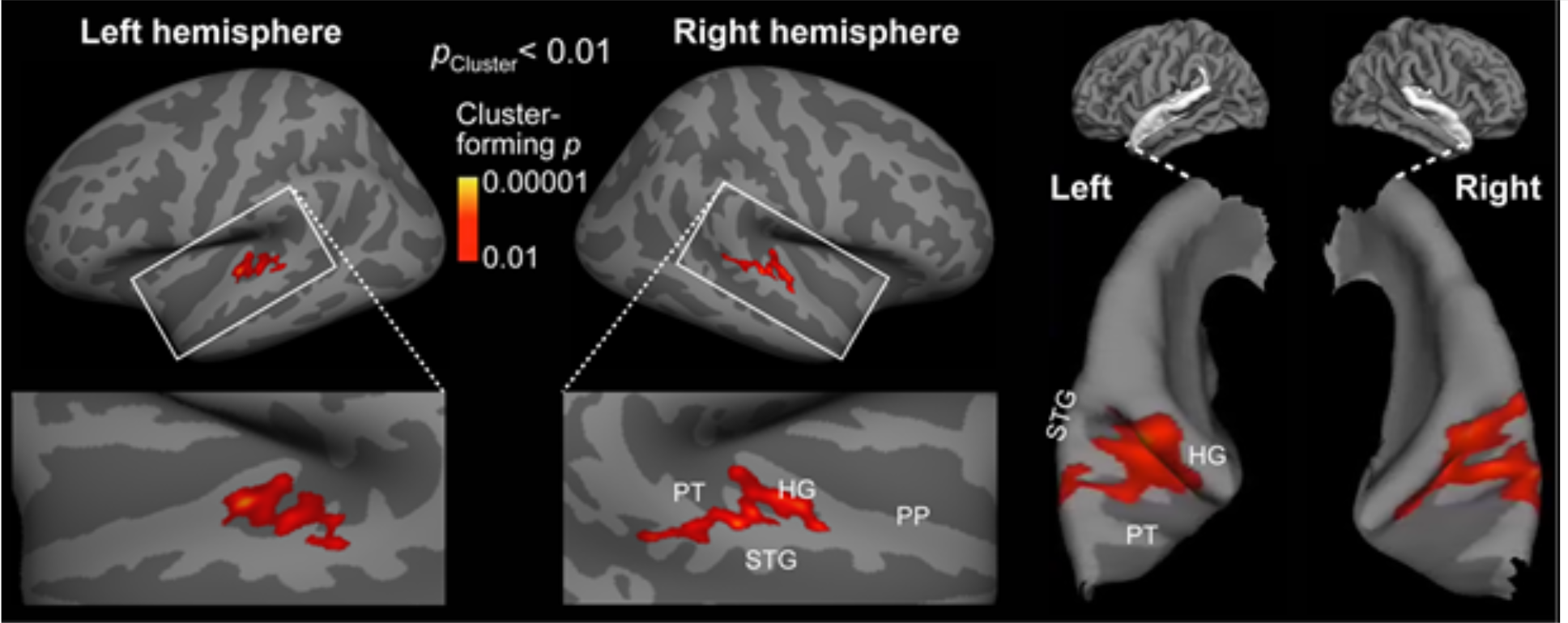
Auditory cortex fMRI activations representing intensity-independent distance for stimuli from straight ahead of the listener. The contrast between fMRI activations to sounds varying in distance vs. those varying in intensity only is shown in inflated left and right hemisphere cortex representations. The centroid of activations was located in posterior non-primary AC areas, overlapping the putative “where” processing stream. The right panel shows the same surface-based activation estimates rendered atop patches of pial surface curvature representations.

In an additional hypothesis-based fMRI analysis (Fig. 4), a region of interest (ROI) was defined in each hemisphere by combining two anatomical FreeSurfer standard-space labels, which encompass PT and posterior aspect of STG. Exactly the same ROIs was used in our previous study that investigated fMRI activations to distance cues from the side of the head (Kopco et al., 2012). Consistent with the whole-brain mapping results (Fig. 3), blood oxygen level dependent (BOLD) percentage signal changes were significantly stronger during varying distance than varying intensity conditions both in the left-hemisphere (*t*_Initial_(10)=2.5, *p*_Corrected_<0.05) and right-hemisphere (*t*_Initial_(10)=3.2, *p*_Corrected_<0.01) ROIs, as tested with a non-parametric randomization test (Blair and Karniski, 1993; Manly, 1997). The difference between the two hemispheres, however, remained non-significant.

**Figure 4.**
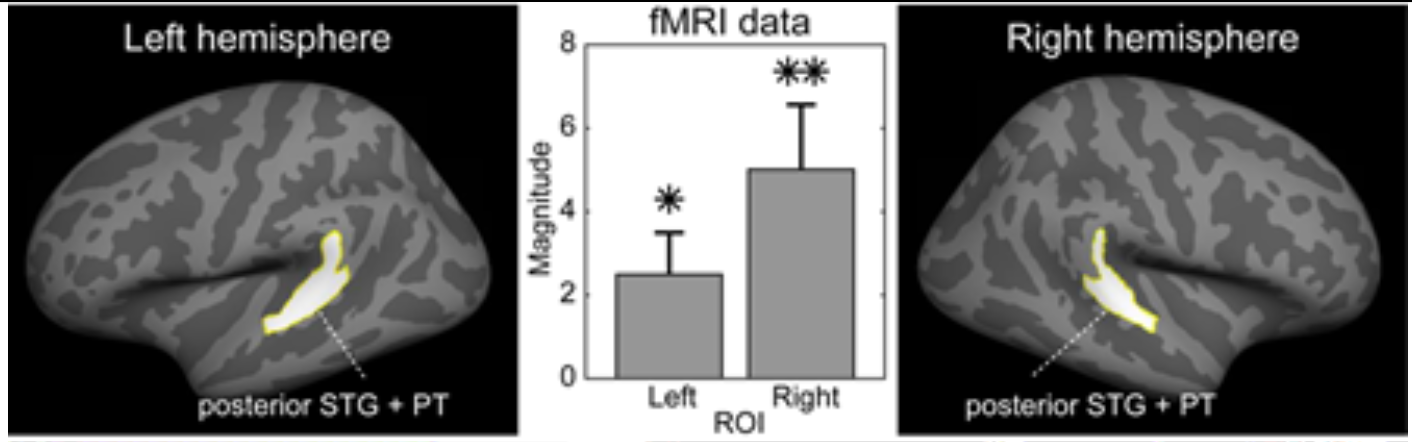
Hypothesis-based region-of-interest (ROI) analysis of distance-cue related posterior non-primary auditory cortex activations. A significant increase of posterior auditory cortex ROI activity was observed during varying distance vs. varying intensity conditions, and more strongly so in the right hemisphere. The values represent contrast effect size values, converted to PSC and normalized by the variance of the PSC. The error bars reflect standard error of mean (SEM). \**p* < 0.05, \*\**p* <0.01, post-hoc corrected based on (Blair and Karniski, 1993; Manly, 1997).

Finally, to enhance comparability to fMRI studies in volume space, we conducted a whole-brain analysis in a 3D standard brain (Fig. 5). In the varying distance vs. varying intensity contrast, a significant activation cluster was identified. The strongest and largest cluster (GRF cluster *p*<0.001, volume 2496 mm^3^, peak voxel [*x, y, z*]_MNI-Talairach_=[64, −37, 7]) was observed in posterior aspects of right AC, which extended from HG to PT and to the posterior crest of STG and the upper bank of superior temporal sulcus (STS). In the left hemisphere, the significant cluster extended from HG to posterior STG and PT (GRF cluster *p*<0.01, volume 1712 mm^3^, peak voxel [*x, y, z*]_MNI-Talairach_ = [−46, −19, 7]).

**Figure 5.**
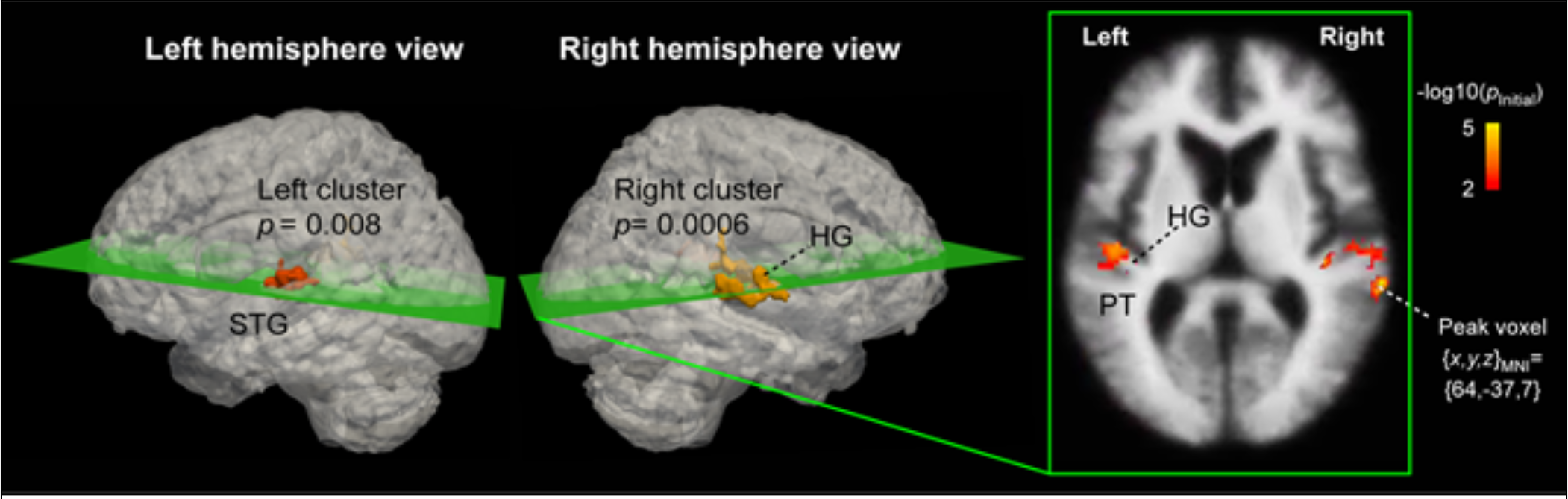
Volume-based fMRI analysis of activations during varying distance vs. intensity. Significant activation clusters, corrected for multiple comparisons based on the GRF theory, extend from HG to PT and posterior aspects of STG in both hemispheres, with the activations being stronger and statistically more powerful in the right than left hemisphere. The left and middle panels show the two significant volume clusters embedded into the respective lateral views of a “glass brain” representation. The right panel depicts the initial cluster-forming functional activation estimates masked in to the significant clusters, as shown in the slice including the largest activation voxel.

## Discussion

We examined the intensity-independent neural representation of auditory distance for frontal sources, for which the representation is not based on ILD. The comparisons of fMRI activations to the Varying-Frontal-Distance vs. Varying-Intensity-at-Fixed-Fronal-Distance conditions revealed, in both hemispheres, a significant activation cluster that was centered in the posterior non-primary ACs. Posterior non-primary areas of human AC have previously been shown to be activated by other auditory-spatial attributes including sound direction changes and auditory motion stimuli (Ahveninen et al., 2014; Griffiths and Warren, 2002; Rauschecker, 2015). Our recent study provided evidence that roughly the same posterior AC areas also contain neuron populations that are sensitive to changes in source distance simulated from the side of the listener’s head, for which distance discrimination can be based on ILD (Kopco et al., 2012). The present results extend these previous studies by providing novel evidence of intensity-independent distance representations that do not involve the ILD. Since ILD is a cue that primarily encodes directional information, these results confirm that the auditory distance area identified in the previous and current studies encodes source distance independent of its direction (or directional cues), even if the distance and direction representations are overlapping.

Many previous fMRI and neurophysiological studies of within and beyond auditory cortex have taken advantage of stimulus-specific adaptation, which results in suppression of responses to repetitive stimuli that fall within the same receptive field (Grill-Spector et al., 2006; Jääskeläinen et al., 2011; Ulanovsky et al., 2003). The idea has been that areas sensitive to a given stimulus feature can be revealed based on a release from this suppression after the presentation of successive stimuli that differ along this feature dimension (Ahveninen et al., 2006; Grill-Spector et al., 2006; Jääskeläinen et al., 2007). However, this strategy can be confounded in auditory distance studies as changes in distance are accompanied by changes in intensity, which in turn results in release from suppression for various feature detectors, in particular when the stimulus is broadband noise containing an abundance on spectro-temporal features. Therefore, the current experiment was designed to contrast conditions with varying distance and received intensity vs. varying only in the received intensity. The observed significant increase in the activation in PT and posterior STG is expected to reflect only representation sensitive to distance, as detectors sensitive to other features are assumed to be equally active in both conditions contrasted here. A supporting fMRI experiment performed in the previous study (Kopco et al., 2012) compared responses to stimuli varying in distance, after normalizing the received intensity, with responses to stimuli at fixed distance and intensity. Widespread and nonspecific activation pattern was observed, confirming that such normalization cannot be easily achieved and that various feature detectors are released from adaptation even for stimuli with “normalized” intensity. Specifically, the current activation pattern shown in Fig. 4 is much less spread into PP and PT regions in both hemispheres than the activation pattern from the previous study (Fig. S3 in Kopco et al., 2012).

The extent of significant activations to Varying Frontal Distance vs. Varying Intensity was slightly wider than in our previous study, in which the distances were simulated from the right side of the listener and activations were only observed in the left hemisphere (Kopco et al., 2012). Also, in the current study, the activations tended to be larger and more posterior in the right than the left hemisphere. Due to the inherent limitations of non-invasive neuroimaging, it is not feasible to discuss whether or not these differences in the size of activated areas reflect the different stimulation directions and whether there are left/right asymmetries in the activations to frontal sources. For example, a likely explanation of the difference between the current and the previous study is in the greater sensitivity offered by the present AB-blocked analysis vs. the randomized trial-design that was used in the main experiment in our previous study. The current AB-blocked design seems to have helped reveal sensitivity to spatial distance cues not only in the posterior non-primary, but also closer to the core areas of auditory cortex involved in the ascending “where” pathway of the primate auditory system (Rauschecker and Tian, 2000). Overall, however, comparison of the current and previous study allows us to conclude that, for frontal sources, distance representation is bilateral (current study), while for sources presented along the interaural axis on the right-hand side, the activation is only observed in the left auditory cortical areas (Kopco et al., 2012).

Consistent with our previous study (Kopco et al., 2012), the centroids of distance-related activations (Varying Frontal Distance > Varying Intensity) were located in superior temporal cortex areas posterior to HG, which have been shown to be associated with other aspects of spatial hearing as well (Ahveninen et al., 2006; Baumgart et al., 1999; Deouell et al., 2007; Warren and Griffiths, 2003; Warren et al., 2002). Interestingly, roughly the same areas showed overlapping activations to individually presented ILD and ITD cues in a recent fMRI study (Higgins et al., 2017). It is thus possible that the posterior AC areas that were activated by frontal distance cues include neuron populations that process representations of the auditory space instead of individual acoustic features only (Higgins et al., 2017; Palomäki et al., 2005), although these representations may not have an orderly topographical organization comparable to cortical representations of the visual space (Salminen et al., 2009; Stecker and Middlebrooks, 2003).

An important question for future studies is whether present activations reflect neuron groups specialized in encoding the DRR (or other reverberation-related) acoustic cue or those representing locations of the acoustic space based acoustic cue combinations (for a review, see (Ahveninen et al., 2014)). The only distance cue independent of overall level changes that is available from the frontal direction is the ratio between direct sounds and reflections, i.e, DRR. Previous psychoacoustic studies suggest that listeners are capable of discriminating source distances based on this cue alone (Kopco and Shinn-Cunningham, 2011). However, in the present Varying Frontal Distance condition, the stimuli contained both DRR and intensity-related distance cues, i.e., all possible available cues. Our previous behavioral studies suggest that in cases like this, auditory distance-discrimination performance is based on an integrated representation of source distance cues instead of any individual distance cue alone (level, ILD, or DRR) (Kopco et al., 2012), consistent with recent fMRI findings regarding horizontal direction cue processing (Higgins et al., 2017). The areas activated more strongly to frontal distance changes than intensity changes could thus involve both DRR specific populations and networks that assemble independent features to a more integrated spatial representation. Our hypothesis for future studies is that neurons sensitive to DRR alone are more prevalent closer to HG and that those sensitive to feature combinations originate further away from the AC core.

The present study concentrated on frontal-distance representation in the near-head range, to be comparable to the lateral-direction distance study of Kopco et al. (2012) in which robust ILD-based distance cues were available only for nearby sources. Therefore, an important question is how these results generalize to larger distances. Since DRR varies with source distance at a constant rate independent of distance (Fig. 1B), it can be expected that, for the frontal-distance judgments based on DRR examined here, the observed distance representation would generalize to larger distances as well. Importantly, DRR and its representation would become dominant also for lateral sources at larger distances, for which ILD becomes unavailable. Therefore, it is expected that the patterns of activation observed here for nearby sources are generalizable to representations of more distant sources as well.

The current behavioral results showed that intensity-independent distance discrimination performance is much better for lateral than for frontal sources. This result is consistent with the hypothesis that ILD contributes to distance judgments for the lateral sources (Kopco et al., 2012). However, it cannot be considered as a conclusive proof of that suggestion, as the DRR cue also varies across a smaller range for the frontal sources than for the lateral sources (in Fig. 1B, frontal DRR varies over approximately 10 dB over the examined range, while the lateral left-ear DRR varies over approximately 20 dB). Thus, it can be expected that frontal distance judgements would be worse than lateral distance judgements even if both of them were based only on DRR (Kopco and Shinn-Cunningham, 2011). In addition, a detailed examination of the dependence of frontal DRR on distance shows that the cue provides very little distance information for the nearest distances examined here (the dashed DRR lines in Fig. 1B are approximately flat for distances 15-25 cm). Despite that, no evidence for a deterioration in discrimination for the nearest distance pairs was observed (thin dashed lines in Fig. 2 are flat). It is likely that other cues, like the acoustic parallax (Kim et al, 2001), supplemented the reverberation-related distance information for these sources. Finally, it is important to note that while it is easy to acoustically compute the DRR of an artificially designed stimulus, is rather difficult to extract it neurally from a sound heard in a real environment as the direct and reverberant portions of the signal overlap in time (Larsen et al, 2008). And, there are many reverberation-related cues that correlate with DRR which might be easier to extract by the brain than DRR. Cues that have been proposed in previous studies are the early-to-late power ratio (Bronkhorst and Houtgast, 1999), the interaural coherence (Bronkhorst, 2001), monaural changes in the spectral centroid or in frequency-to-frequency variability in the signal (Larsen et al., 2008), and amplitude modulation (Kim et al., 2015; Kolarik et al., 2016), all of which vary systematically as a function of distance in reverberant environments. Future fMRI studies can compare neural activation in response to these cues with those in response to varying distance and varying DRR to identify which of them is likely the cue encoded by listeners’ brains when judging distance in reverberation.

## Conclusions

Our results suggest that posterior human auditory cortex areas contain neurons that are sensitive to distance cues like DRR that are independent of intensity and binaural cues relevant for direction hearing. The behavioral experiment further demonstrated that these frontal intensity-independent cues are perceptually relevant for human listeners.

## Acknowledgments

This work was supported by the EU H2020-MSCA-RISE-2015 grant no. 9122, the EU RDP projects TECHNICOM I, ITMS: 26220220182, and TECHNICOM II, ITMS2014+:313011D23, and the SRDA, project APVV-0452-12. The imaging studies were also supported by the NIH grants R01DC017991, R01DC016915, R01DC016765, and R21DC014134. Zoltan Szoplak contributed to the behavioral experiment data collection. The authors declare no competing interests.

